# A designer FG-Nup that reconstitutes the selective transport barrier of the Nuclear Pore Complex

**DOI:** 10.1101/2020.02.04.933994

**Authors:** Alessio Fragasso, Hendrik W. de Vries, Eli O. van der Sluis, Erik van der Giessen, Patrick R. Onck, Cees Dekker

## Abstract

Nuclear Pore Complexes (NPCs) regulate bidirectional transport between the nucleus and the cytoplasm. Intrinsically disordered FG-Nups line the NPC lumen and form a selective barrier, where transport of most proteins is inhibited whereas specific transporter proteins freely pass. The mechanism underlying selective transport through the NPC is still debated. Here, we reconstitute the selective behaviour of the NPC bottom-up by introducing a rationally designed artificial FG-Nup that mimics natural Nups. Using QCM-D, we measure a strong affinity of the artificial FG-Nup brushes to the transport receptor Kap95, whereas no binding occurs to cytosolic proteins such as BSA. Solid-state nanopores with the artificial FG-Nups lining their inner walls support fast translocation of Kap95 while blocking BSA, thus demonstrating selectivity. Coarse-grained molecular dynamics simulations highlight the formation of a selective meshwork with densities comparable to native NPCs. Our findings show that simple design rules can recapitulate the selective behaviour of native FG-Nups and demonstrate that no specific spacer sequence nor a spatial segregation of different FG-motif types are needed to create functional NPCs.

## Introduction

Nucleocytoplasmic transport is orchestrated by the Nuclear Pore Complex (NPC), which imparts a selective barrier to biomolecules^1,2^. The NPC is a large eightfold-symmetric protein complex (with a size of ∼52 MDa in yeast and ∼112 MDa in vertebrates) that is embedded within the nuclear envelope and comprises ∼30 different types of Nucleoporins (‘Nups’)^3,4^. Intrinsically disordered proteins, termed FG-Nups, line the central channel of the NPC. FG-Nups are characterized by the presence of phenylalanine-glycine (FG) repeats separated by spacer sequences^5^ and they are highly conserved throughout species^6^. FG-Nups carry out a dual function: By forming a dense barrier (100-200 mg/mL) within the NPC lumen, they allow passage of molecules in a size-selective manner^7–10^. Small molecules can freely diffuse through, whereas larger particles are generally excluded^11^. At the same time, FG-Nups mediate the transport of large NTR-bound (Nuclear Transport Receptor) cargoes across the NPC through transient hydrophobic interactions between FG repeats and hydrophobic pockets on the convex side of NTRs^12^. Various models have been developed in order to connect the physical properties of FG-Nups to the size-selective properties of the NPC central channel, e.g. the ‘virtual-gate’^13^, ‘selective phase’^14,15^, ‘reduction of dimensionality’^16^, ‘kap-centric’^17–19^, ‘polymer brush’^20^, and ‘forest’^5^ models. As is evident from the multitude of transport models, no consensus on the NPC transport mechanisms has yet been reached.

The NPC is highly complex in its architecture and dynamics, being constituted by many different Nups that simultaneously interact with multiple transiting cargoes and NTRs. In fact, translocating cargoes may amount to almost half of the mass of the central channel, so they may be considered an intrinsic part of the NPC^3^. These NPC properties complicate in-vivo studies^3,21–23^, for which it is very challenging to identify contributions coming from individual FG-Nups^24,25^. On the other hand, in-vitro approaches to study nucleocytoplasmic transport using biomimetic NPC systems^26–33^ have thus far been limited to single native FG-Nups and mutations thereof, attempting to understand the physical behaviour of FG-Nups and their interactions with NTRs. The reliance on a few selected Nups from yeast or humans in these studies with sequences that evolved over time in different ways for each of these specific organisms makes it difficult to pinpoint the essential and minimal properties that provide FG-Nups with their specific selective functionality.

Here, we describe a bottom-up approach to studying nuclear transport selectivity, where we rationally design, synthesize, and assess artificial FG-Nups with user-defined properties that are set by an amino acid sequence that is chosen by the user. With this approach we address the question: can we build a synthetic protein that mimics the selective behaviour of native FG-Nups? By combining experiments and coarse-grained molecular dynamics simulations, we illustrate the design and synthesis of an artificial 311-residue long FG-Nup, which we coin ‘NupX’, and characterize its selective behaviour with respect to Kap95 (a well-characterized NTR from yeast, 95 kDa), Bovine Serum Albumine (BSA, 66 kDa), and tCherry (tetramer of mCherry, 104 kDa). Firstly, we explore the interactions between Kap95 and NupX brushes using quartz crystal microbalance with dissipation monitoring (QCM-D), finding that NupX brushes bind Kap95 with a binding affinity comparable with that of native FG-Nups, while showing no binding to BSA or tCherry. We confirm this finding by calculating the potential of mean force (PMF) associated with the entry of Kap95 or an inert cargo into NupX brushes. Secondly, we explore the transport properties of NupX-functionalized solid-state nanopores and show that NupX-lined pores constitute a selective barrier. Similar to FG-Nups previously studied with the same technique^28,31^, the NPC-mimicking nanopores allow fast and efficient passage of Kap95 molecules, while blocking transport of BSA. Coarse-grained MD simulations of NupX-functionalized nanopores highlight the formation of a dense FG-rich meshwork with similar protein densities as in native NPCs, which excludes tCherry but allows entry and passage of Kap95.

The current work provides the proof of concept that a designer FG-Nup can reconstitute NPC-like selectivity, and the results show that no specific spacer sequence nor a spatial segregation of different FG-motifs (as observed in recent work^3,34^) are required for achieving size-selectivity. This work lays the foundation for multiple future directions in follow-up work as the approach opens the route to systematically study the essential microscopic motifs that underlie the unique selectivity of NPCs.

## Results

### Design of the synthetic NupX

In the design of our synthetic NupX protein, we aim to reconstitute nuclear transport selectivity while operating under a minimal set of simple design rules. The design procedure that we outline below uses the following four rules: i) we design a protein that incorporates the physical properties of GLFG-Nups, ii) it comprises two parts, with a cohesive domain at one end and a repulsive domain at the other end, where each domain is characterized by the ratio C/H of the number of charged and the number of hydrophobic residues, iii) FG and GLFG motifs are present in an alternating and uniformly spaced fashion within the protein’s cohesive domain, and iv) the protein is intrinsically disordered throughout its full length, similar to native FG-Nups.

We implemented our design rules in a step-wise design process as follows: First, we selected and analysed an appropriate set of native FG-Nups (design rule i), namely GLFG-Nups, which differ from FxFG-Nups in terms of the type of FG repeats and the properties of the spacer regions^11^. The emphasis on GLFG-Nups follows from their localization in the central channel^3^ of the yeast NPC (Figure 1a), where they strongly contribute to the nuclear transport selectivity. Indeed, a small subset of GLFG-Nups (e.g., either Nup100 or Nup116 in combination with Nup145N) was shown to be essential and sufficient for cell viability^21,35^. To derive the amino acid content of NupX, we therefore characterized the archetypical GLFG-Nup sequence by determining the amino acid content of the disordered regions of Nup49, Nup57, Nup145N, Nup116, and Nup100 from yeast. Of these, the most essential GLFG-Nups (i.e. Nup100, Nup116, and Nup145N) comprise a collapsed domain with a low C/H-ratio and abundance of FG/GLFG repeats, and an extended domain with a high C/H-ratio and absence of FG repeats (Figures 1b,c)^5^. The division into two domains of these essential GLFG-Nups led us to phrase design rule ii in our design process of NupX, with each domain comprising ∼150 amino acid residues (see Figures 1b,c).

**Figure 1:**
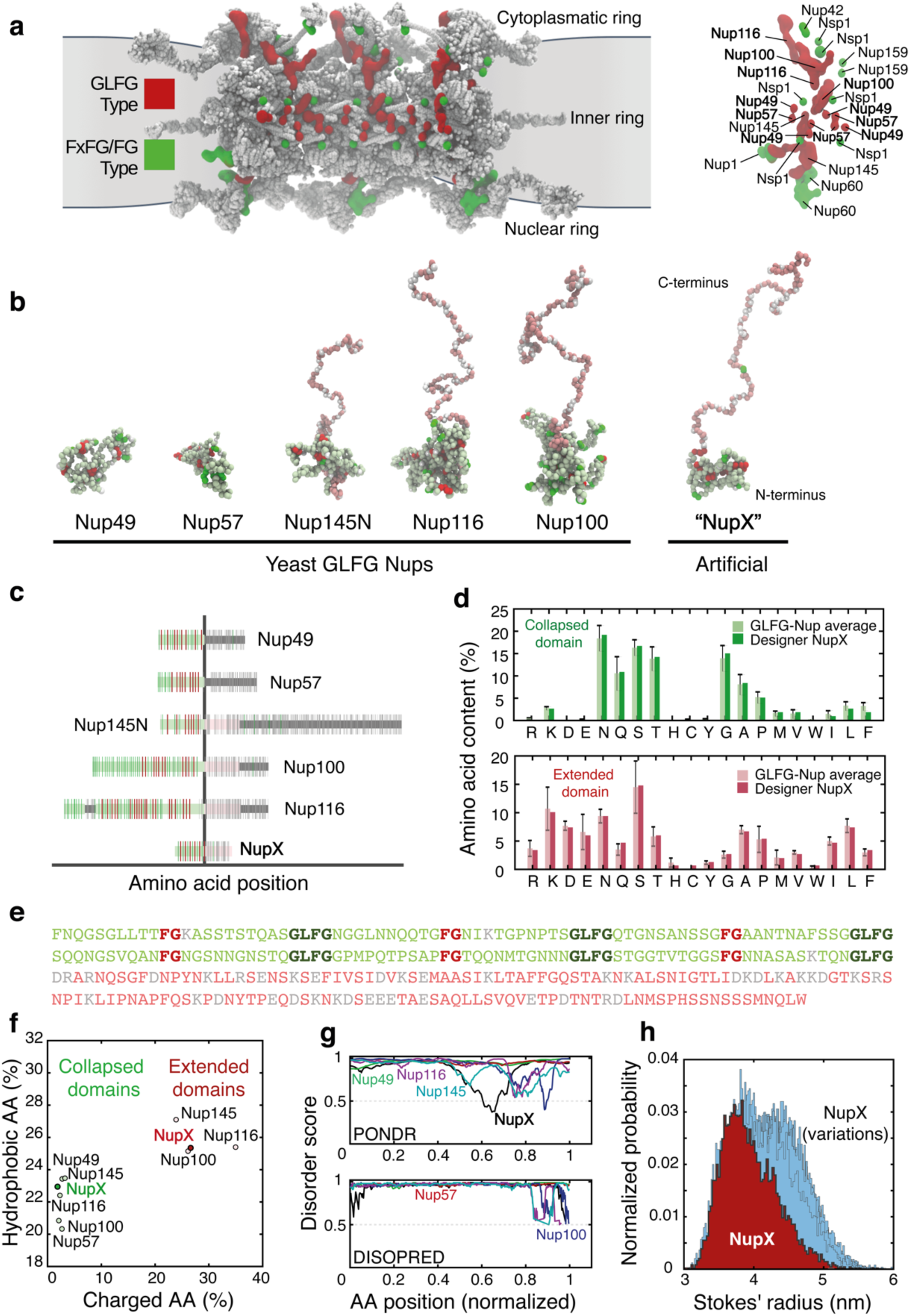
De novo design of an artificial FG-Nup. **a**, Left: Frontal view on three of the eight spokes of the yeast NPC that shows how the GLFG-Nups (red) are predominantly anchored in the inner ring, as opposed to the FxFG/FG-Nups (green), which are anchored towards the cytoplasmic or nuclear rings. Right: highlight of the anchoring points of individual Nups in a single spoke of the NPC. The GLFG-Nups Nup100, Nup116, Nup49, and Nup57 contribute strongly to the permeability barrier of the NPC^3^, where Nup100 and Nup116 are known to be indispensable for NPC viability^21,35^. **b**, Simulation snapshots of isolated native yeast GLFG-Nups at one amino acid resolution. The conformations of Nup145N, Nup116, Nup100 highlight a ‘bimodality’ of the Nups^5^, with a collapsed (light green) and extended (pink) domain. FG repeats, GLFG repeats, and charged residues are displayed in green, red, and white, respectively. NupX adopts the same bimodal conformations as the essential GLFG-Nups Nup100 and Nup116. **c**, Comparison of the full-length sequences between yeast GLFG-Nups and NupX. Sequence highlights follow the same colour-scheme as in panel b, folded domains are indicated in dark-grey. **d**, Amino acid contents of yeast GLFG-Nups (averaged) and NupX for the collapsed (top panel) and extended (bottom panel) domains. FG and GLFG-motifs were excluded from this analysis. The amino acid sequence of NupX is based on the average composition of yeast GLFG-Nup domains. **e**, Designed sequence of NupX, following the same color scheme as figures b,c. FG and GLFG repeats are spaced by 10 residues in the cohesive low C/H-ratio domain. **f**, Charge-and-hydrophobicity plot of yeast GLFG-Nup domains as compared to domains of NupX. For both the collapsed and extended domains, the charged and hydrophobic amino acid contents of NupX agree with the properties of individual GLFG-Nups. **g**, Disorder prediction scores for the unfolded domains of GLFG-Nups and full-length NupX from two different predictors (see Materials and Methods). Disorder prediction scores higher than 0.5 (dashed line) count as fully ‘disordered’. **h**, Distribution of Stokes radii from 10 μs of coarse-grained molecular dynamics simulations for both NupX (red) and 25 design variations (light blue). NupX is, on average, slightly more compacted as compared to other design variants.

Assigning the amino acid (AA) content to NupX, as derived from the sequence information of the GLFG-Nups, was performed separately for the two domains: we computed the cumulative amino acid contents (excluding FG and GLFG motifs) for both the collapsed domains of all five GLFG-Nups, and for the extended domains of Nup100, Nup116 and Nup145N (design rule ii). Upon normalizing for the total length of the collapsed or extended domains of all native GLFG-Nups, this analysis resulted in the distributions presented in Figure 1d, plotted separately for the collapsed (light green, top) and the extended (light red, bottom) domains. Based on these histograms, we assigned amino acids to the collapsed and extended domains of NupX separately. Following design rule iii, we then placed FG and GLFG repeats in the collapsed domain with a fixed spacer length of 10 AAs. This value was chosen based on the spacer length of ∼5-15 AAs in native GLFG-Nups. An analysis of the charged and hydrophobic amino acid content of the domains of NupX and native GLFG-Nups shows that the assigned sequence properties are indeed reproduced by our design method (Figure 1f). Finally, the sequences of the collapsed and extended domains of NupX were repetitively shuffled (except for the FG and GLFG motifs that we kept fixed) until a desirable level of disorder was achieved (design rule iv), as predicted by PONDR^36^ and DISOPRED^37,38^ (Figure 1g). This resulted in the NupX sequence shown in Figure 1e. Whereas PONDR predicts one short folded segment between residues 189 and 209 (normalized position of 0.65 in Figure 1g), additional structure prediction^39^ (Materials and Methods) did not yield any high-confidence folded structures for this segment.

To assess the robustness of our design procedure, we tested how permutations of the NupX sequence (which shuffle amino acids while retaining the FG/GLFG sequences and the definition of both domains) affect the Stokes radius *R*_*S*_, as computed from 1-bead-per-amino-acid MD-simulations developed for intrinsically disordered proteins (Figure 1h, see Materials and Methods). We found that over 25 different designs for NupX (Table S2) yielded an average *R*_*S*_ of 4.2 ± 0.2 nm (errors are S.D.). This is close to the simulated (3.9 ± 0.4 nm) and measured (3.7 ± 1.1 nm by DLS, Table S1) *R*_*S*_ value of the NupX protein design (Figure 1e).

Summing up, using a minimal set of rules, we designed a NupX protein that incorporates the average properties that characterize GLFG-Nups^5,11^. Moreover, by creating 25 different designs that all showed similar behaviour in our simulations, we showed that the physical properties such as the Stokes radius and the division of NupX into a cohesive and repulsive domain are recovered in a reliable way.

### QCM-D experiments and MD simulations show selective binding of Kap95 to NupX brushes

To assess the interaction between NupX and Kap95, we employed a Quartz-Crystal-Microbalance with Dissipation monitoring (QCM-D), with gold-coated quartz chips and phosphate-buffered saline (PBS, pH 7.4) as running buffer, unless stated otherwise. First, C-terminus-thiolated NupX molecules were injected into the chamber at a constant flow-rate (20 μL/min) where they chemically reacted with the gold surface (Figure 2a). Binding of NupX to the gold surface could be monitored in real-time by measuring the shift in resonance frequency Δ*f* of the quartz chip. Data were fitted by the Voigt-Voinova viscoelastic model^40^ to extract the areal mass density of the deposited film (see Materials and Methods and Table S4), which, as widely reported for QCM-D experiments^41–43^, measures both the protein mass as well as the hydrodynamically coupled water to it. Assuming 35% of the sensed film to be constituted by protein mass^43^, with the remaining 65% being constituted by coupled water, we estimated an average grafting distance *d*_*g*_ between adjacent NupX proteins of *d*_*g*_∼5.6 nm, assuming a triangular lattice (since an equilateral triangulated (hexagonal) lattice is the densest type of packing that can be described by a unique length scale that sets the grafting density). After the Nup-layer was formed, a 1-mercapto-11-undecylte-tra(ethyleneglycol) molecule, which is expected to form a ∼2 nm thin passivating film^17^, was added to passivate any remaining bare gold that was exposed in between NupX molecules, which minimizes unintentional interactions between Kap95 and gold (Figure S3)^17,18,45^.

**Figure 2:**
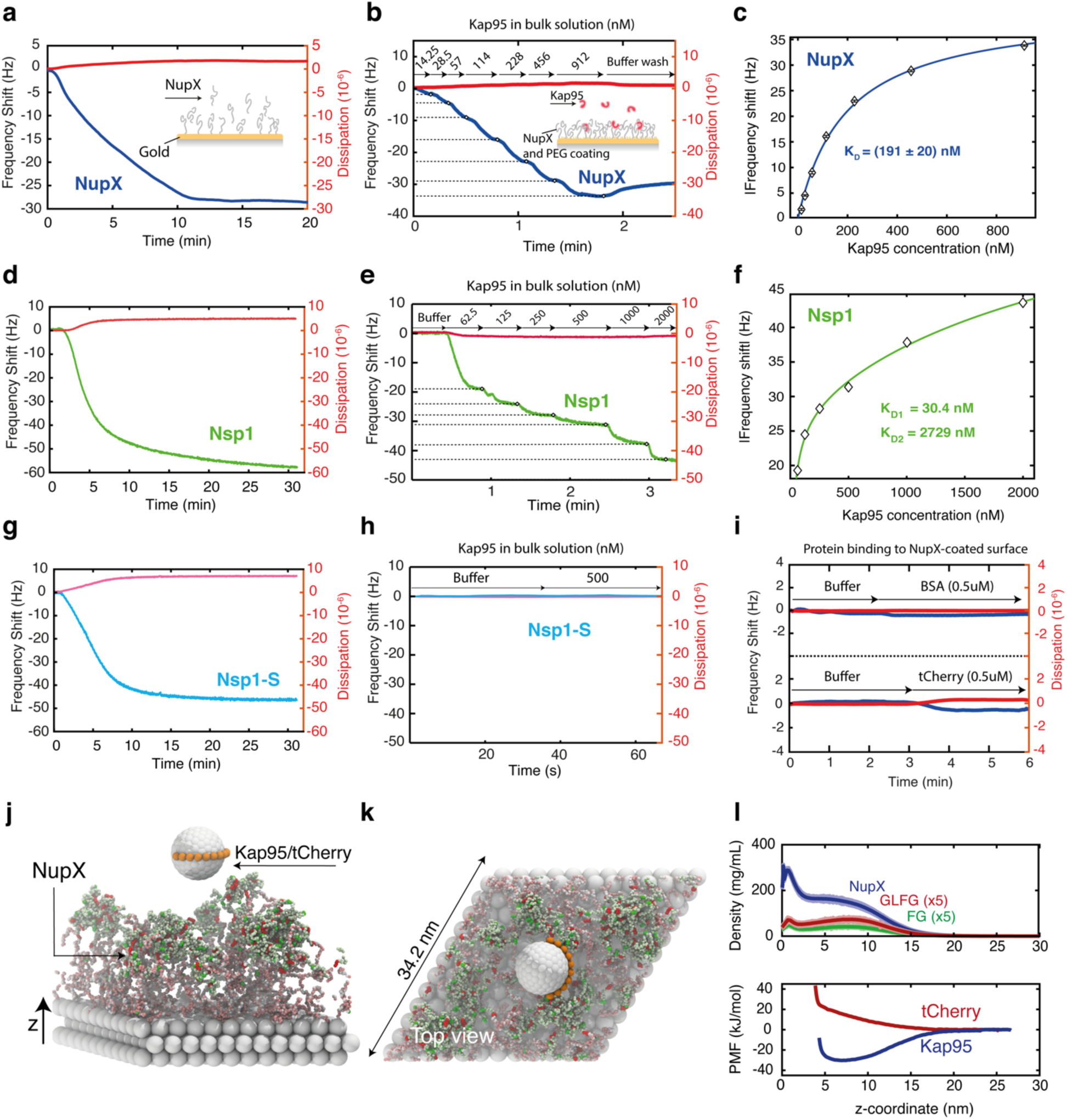
Binding affinity of Kap95 to NupX, Nsp1 and Nsp1-S brushes, using QCM-D and MD simulations. **a,d,g**, Coating of gold surface with NupX, Nsp1, and Nsp1-S proteins, respectively, at 1 µM protein concentration. **b,e,h**, Change in frequency shift upon titration of Kap95 (with concentration in the range ∼10-2000 nM) on NupX, Nsp1, and Nsp1-S coated surfaces. Numbers indicate the concentration in nM of Kap95 for each titration step. Large changes in frequency shift are observed for NupX and Nsp1, whereas no detectable shift is measured for Nsp1-S. **c,f**, Frequency shift upon titration of Kap95, with fits by single (blue curve in b, for NupX) or two-component (green curve in **e**, for Nsp1) Langmuir isotherms. **i**, Frequency shift upon adsorption of BSA (top) and tCherry (bottom) onto the NupX-coated sensor. **j**, Side-view snapshot of the umbrella sampling simulation setup, where a model Kap95 particle (8.5 nm diameter) or tCherry-sized particle (7.5 nm diameter, not shown) is restrained along different z-coordinates. Scaffold beads are shown in grey, NupX proteins follow the same color scheme as presented in Figure 1b. **k**, Top-view of the periodic triclinic simulation box. **l**, Top panel: Time and laterally averaged protein density distributions of NupX brushes and for the two different types of FG motifs present inside the NupX proteins. The density profiles of the GLFG and FG motifs within the NupX brush are multiplied by 5 for clarity. Light shades indicate the standard deviation in the time-averaged density profile. A high-density region (up to 300 mg/mL) forms near the attachment sites (*z* = 0 to 2.5 nm). Further away from the scaffold, the protein density remains at a constant value of ∼ 170 mg/mL up to a distance of ∼8 nm, after which it decays. FG and GLFG motifs predominantly localize near the transitioning point (8 nm). Bottom: Free-energy profiles (PMF-curves) of the center of mass of the model Kap95 and tCherry-sized inert particle along the *z*-coordinate, where *z* = 0 coincides with the substrate. The difference in sign between the PMF-curves of both particles indicates a strong preferential absorption of the model Kap95 to NupX brushes and a repulsive interaction with tCherry.

After thus setting up a NupX-coated layer, we flushed in Kap95 at stepwise increasing concentrations (∼10-1000 nM, Figure 2b) and monitored binding to the NupX-coated surface. We observed a clear concentration-dependent absorption of Kap95 molecules to the NupX brush. We fitted a single Langmuir isotherm to the saturation points, see Figure 2c, and thus obtained a binding affinity of *K*_*d*_ = 191 ± 20 nM (error is S.D.). This value compares very well to *K*_*d*_ values for the binding between Kap95 and native FG-Nups as found with similar techniques, which typically range between 100 and 500 nM^18,27^. For reference, we similarly measured the affinity of Kap95 to Nsp1, a native FG-Nup from yeast, as well as to Nsp1-S, a Nsp1-mutant where the hydrophobic amino acids F, I, L, V are replaced by the hydrophilic amino acid Serine (S), which serves as a negative control since it is expected to not bind Kap95 due to the lack of FG repeats^14,15^ (Figure 2d,g). Upon binding of Nsp1 and Nsp1-S to the gold surface under the same conditions as for NupX, we measured a *d*_*g*_∼5.8 nm and *d*_*g*_∼5.4 nm for Nsp1 and Nsp1-S, respectively, gratifyingly close to the value for NupX. Consistent with previous studies^27,45^, we found that Kap95 exhibits a concentration-dependent absorption to Nsp1 brushes (Figure 2e), whereas we did not observe any detectable interaction between Kap95 and Nsp1-S (Figure 2h). The latter is consistent with the lack of FG repeats in the Nsp1-S sequence which makes the Nsp1-S film devoid of binding sites for Kap95. For the Nsp1-Kap95 interaction, we found dissociation constants of *K*_*d*,1_ = 30 ± 25 nM and *K*_*d*,2_ = 2729 ± 626 nM upon fitting a two-component Langmuir isotherm (Figure 2f) to the saturation points from the frequency diagram of Figure 2e. Using a single isotherm yielded a poor fit to the data (Figure S5) with *K*_*d*_ = 94 ± 36 nM, of the same order as the *K*_*d*_ for NupX. That the Kap95 to NupX and Nsp1(-S) binding constants were found using different fits (single component vs. two-component Langmuir isotherms, respectively) can be attributed to differences between NupX and Nsp1 in terms of their respective ratios of *d*_*g*_ and *R*_*S*_. In previous work^18,46^, sparsely grafted FG domains (*d*_*g*_ > *R*_*S*_) revealed a single type of binding when interacting with Impβ (Kap95 human homolog) molecules, as opposed to closely-packed FG domains (*d*_*g*_ < *R*_*S*_) that instead showed two binding modes. Following this reasoning, the use of a two-component isotherm for the Nsp1(-S) brushes would be consistent with these and other previous studies^17,18^, since *R*_*S*_ (7.9 ± 2.0 nm for Nsp1, 6.8 ± 1.6 nm for Nsp1-S; errors are S.D.; table S1) is larger than *d*_*g*_ (∼5.4-5.8 nm) for these proteins. For NupX, the opposite holds, since the value of *R*_*S*_ (3.7 ± 1.1 nm S.D. table, S1) is smaller than *d*_*g*_ (∼5.6 nm) and therefore consistent with the observed type of binding.

Flushing a fresh buffer solution without protein after Kap95 titration induced a relatively slow dissociation of Kap95 from both the NupX and Nsp1 brushes, similar to other work^17,18^ Absorbed molecules could be completely removed upon flushing 0.2 M NaOH however (data not shown). Finally, we investigated whether the inert molecules BSA and tCherry could bind to the NupX brush. Upon flushing in 500 nM of BSA (Figure 2i, top) or 500 nM tCherry (Figure 2i, bottom), we observed only a negligible change (< 1 Hz) in the resonance frequency, indicating that the NupX brush efficiently excludes these inert molecules. Importantly, the data thus shows that the NupX selectively interacts with Kap95.

In order to study the morphology and physical properties of NupX brushes at the microscopic level, we employed coarse-grained molecular dynamics (MD) simulations (see Materials and Methods), which resolved the density distribution within the NupX brush layer and the preferential absorption of Kap95 over inert molecules such as tCherry. 36 NupX proteins were tethered on a triangular lattice with a fixed spacing of 5.7 nm (Figure 2j,k). Averaged over a simulation time of 3 μs, we found that the NupX brushes form meshwork with densities ranging from ∼300 mg/mL near the substrate to ∼170 mg/mL throughout the central region of the brush (Figure 2l, top panel). The interface near the free surface of the brush contains the highest relative concentration of FG and GLFG motifs (see Figure 2l, top panel). Notably, the protein density throughout the brush is of the same order of magnitude as the density obtained in simulations of the yeast NPC^47^.

To assess the size-selective properties of the NupX brushes, we performed umbrella sampling simulations of the absorption of Kap95 and tCherry to NupX brushes (Materials and Methods). We modelled Kap95 (Figure S14) as an 8.5 nm sized sterically repulsive (i.e., modelling only repulsive, excluded volume interactions) particle with 10 hydrophobic binding sites^8,31,48,49^ and a total charge density similar to that of Kap95 (−43e). tCherry was modelled as a sterically inert spherical particle of 7.5 nm diameter^10^. We obtained potential-of-mean-force (PMF) curves associated with the absorption of Kap95 and tCherry particles by means of the weighted histogram analysis method (WHAM)^50^. We found that a significant (−30.3 kJ/mol) negative free energy is associated with the entry of Kap95 in the NupX brush, as is visible in Figure 2l (bottom panel), which corresponds to a considerable (∼5 μM) binding affinity. By contrast, the PMF curve of tCherry steeply increased when the protein entered into the NupX meshwork, showing that absorption of non-specific proteins of comparable size as Kap95 will not occur. While the large free energy differences between Kap95 and tCherry adsorption thus qualitatively support the experimental findings, the binding strength of Kap95 to NupX-brushes is quantitatively weaker in our simulations than observed in our experiments. We note however, that a precise numerical correspondence in binding affinity should not be expected given the simplicity of our Kap95 and tCherry models, which were not quantitatively parametrized against binding to NupX.

### Single-molecule translocation experiments with NupX-coated nanopores demonstrate selectivity

In order to test whether our synthetic FG-Nup do indeed form a transport barrier that mimics the size-selective properties of the NPC, we performed electrophysiological experiments on biomimetic nanopores^28,31^. These NPC mimics were built by tethering NupX proteins to the inner walls of a solid-state SiN_x_ nanopore^51^ using Self-Assembled-Monolayer (SAM) chemistry (details in Materials and Methods). Solid-state nanopores of 10-60 nm in diameter were fabricated onto a glass-supported^52^ SiN_x_ free-standing membrane by means of TEM drilling. A buffer with 150 mM KCl, 10 mM Tris, 1 mM EDTA, at pH 7.5 was used to measure the ionic conductance through the pores, while retaining near-physiological conditions. Coating bare SiN_x_ pores with NupX yielded a significant decrease in conductance (e.g. ∼50% for ∼30 nm diameter SiN pores) of the bare-pore values, as estimated by measuring the through-pore ionic current before and after the functionalization (Figure S7). Additionally, the current-voltage characteristic in the ±200 mV range (Figure S7) is linear both for the bare and NupX-coated pores, indicating that the NupX meshwork is not affected by the applied electric field at the 100 mV operating bias. To obtain more information on the NupX-coating process of our SiN_x_ pores, we repeated the same functionalization procedure on SiN-coated quartz chips, while monitoring the shift in resonance frequency using QCM-D (Figure S2), while keeping the protein concentration and incubation time the same as for the coating of the SiN_x_ nanopores for consistency. From this experiment, we estimate an average grafting distance of ∼6.2 nm between adjacent NupX molecules. Measurements of the ionic current through NupX-coated pores revealed a higher 1/f noise in the current (Figure S9) compared to bare pores, which we attribute to random conformational fluctuations of the Nups within the pore volume and access region^53,54^, similar to findings from previous studies on biomimetic nanopores^28,31^.

To test the selective behaviour of the biomimetic nanopore, we measured translocation rates of Kap95 and BSA through bare pores of ∼30-35 nm in diameter (Figure 3a). Figure 3c shows examples of raw traces recorded for a 30 nm pore under 100 mV applied bias, when either only buffer (top), 450 nM Kap95 (middle), or 2.8 µM BSA (bottom) were added to the cis-chamber. As expected, we observed transient dips in the current through the bare pore upon injection of the proteins, which we attribute to single-molecule translocations of the analyte molecules. As is typical in nanopore experiments, translocation events yield current blockades with a characteristic amplitude and dwell time, where the former relates to the size of the molecule occupying the pore and the latter generally depends on specific interaction between the translocating molecule and the pore wall^55^. Next, we repeated the experiment under identical conditions on the same pore after coating with NupX took place (Figure 3b). Examples of typical raw traces are shown in Figure 3d. Strikingly, Kap95 molecules could still translocate efficiently through the NupX-coated pore, whereas BSA molecules were practically blocked from transport.

**Figure 3.**
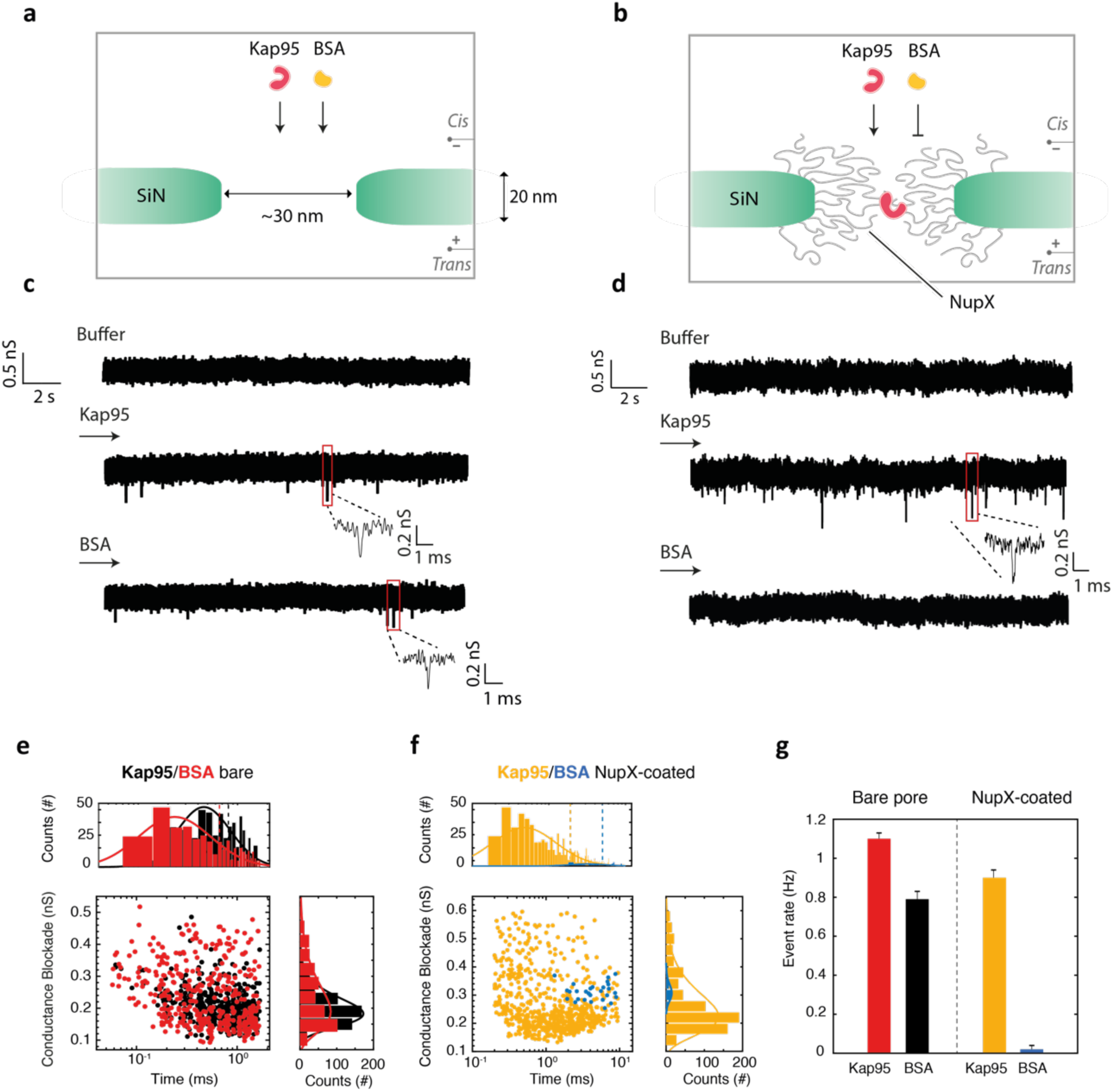
Electrical measurements on NupX-coated solid-state nanopores. **a-b**, Schematic of the nanopore system before (a) and after (b) NupX functionalization. **c-d**, Examples of raw current traces through bare (c) and NupX-coated (d) pores, recorded under 100 mV applied bias for different analyte conditions. Current traces are recorded in the presence of buffer only (top), upon addition of 450 nM Kap95 (middle), and 2.8 uM BSA (bottom). Traces were filtered at 5 kHz. **e**, Scatter plot showing conductance blockades and dwell time distributions of translocation events of the analytes Kap95 (N=506) and BSA (N=387) through a bare 30 nm pore, recorded over the same time interval. **f**, Scatter plot showing conductance blockades and dwell time distributions of translocation events of the analytes Kap95 (N=686) and BSA (N=28) through a NupX-coated 30 nm pore, recorded over the same time interval. Top and right panels in **e** and **f** show lognormal fits to the distribution of dwell times and conductance blockades, respectively. Dashed vertical lines in top panels indicate the mean values for the dwell time distributions. **g**, Event rate of translocations through bare (left) and NupX-coated (right) pore for Kap95 and BSA.

Figures 3e-f show scatter plots of the event distributions, where the conductance blockade is plotted against dwell time for all translocation events. For the bare pore, we observe similar average amplitudes of 0.24 ± 0.09 nS and 0.20 ± 0.05 nS (errors are S.D.) for BSA and Kap95, respectively. For the NupX-coated pore, we found slightly larger but again mutually similar event amplitudes of 0.31 ± 0.03 nS and 0.27 ± 0.03 nS for BSA and Kap95, respectively. We found comparable translocation times through the bare pore of 0.66 ± 0.03 ms and 0.81 ± 0.02 ms (errors are s.e.m) for BSA and Kap95, respectively. For the coated pore, however, we measured longer dwell times of 5.0 ± 0.5 ms and 1.9 ± 0.1 ms for BSA and Kap95, respectively, which indicates that the presence of the NupX molecules in the pore significantly slows down the translocation process of the passing molecules. Repeating the same experiment on a larger 60 nm NupX-coated pore (Fig. S8) yielded selective pores with faster translocations for both Kap95 (0.65±0.05 ms) and BSA (1.6±1.3 ms), consistent with the presence of an open central channel. Smaller pores (<25 nm) did not result in any detectable signal for either Kap95 or BSA (data not shown), due to the poor signal-to-noise ratio attainable at such low conductances.

Most importantly, these data clearly show selectivity of the biomimetic pores. Figure 3g compares the event rate of translocations for Kap95 and BSA through bare and NupX-coated pores under 100 mV applied bias. Event rates were 0.79 ± 0.04 Hz and 1.10 ±0.04 Hz (N=3 different nanopores; errors are s.d.) for BSA and Kap95 through the bare pore, respectively, whereas upon coating the pore with NupX, the event rates changed to 0.02 ± 0.02 Hz and 0.90 ± 0.04 Hz (N=3 different nanopores) for BSA and Kap95, respectively. The sharp decrease in event rate for BSA upon NupX coating of the pores indicates that BSA molecules are strongly hindered by the NupX meshwork formed inside the pore. In contrast, the transport rate of Kap95 through the coated pore is nearly unaffected when compared to the bare pore. From these experiments, we conclude that the user-defined NupX does impart a selective barrier, very similar to native FG-Nups^26,28,31^, by allowing efficient transportation of Kap95 while hindering the passage of BSA.

### MD simulations of NupX-lined nanopores reveal their protein distribution and selectivity

We used coarse-grained MD simulations (Materials and Methods) to understand the size-selective properties of NupX-lined nanopores as obtained in our experiments. The 20 nm height of these nanopores is the same as the SiN_x_ membrane thickness, while we vary the diameter from 15 to 70 nm. Multiple copies of NupX are tethered to the nanopore lumen by their C-terminal domain in an equilateral triangular lattice with a spacing of 5.5 nm, in accordance with estimates initially obtained from the QCM-D experiments (Figure 4a, Materials and Methods, Figure. S12). Based on 6 μs of coarse-grained MD simulations, we obtained the protein density distribution in the (*r, z*)-plane (averaged over time and angle *θ*) within a NupX-lined nanopore of 30 nm in diameter (Figure 4b), similar in size as the translocation experiments.

**Figure 4:**
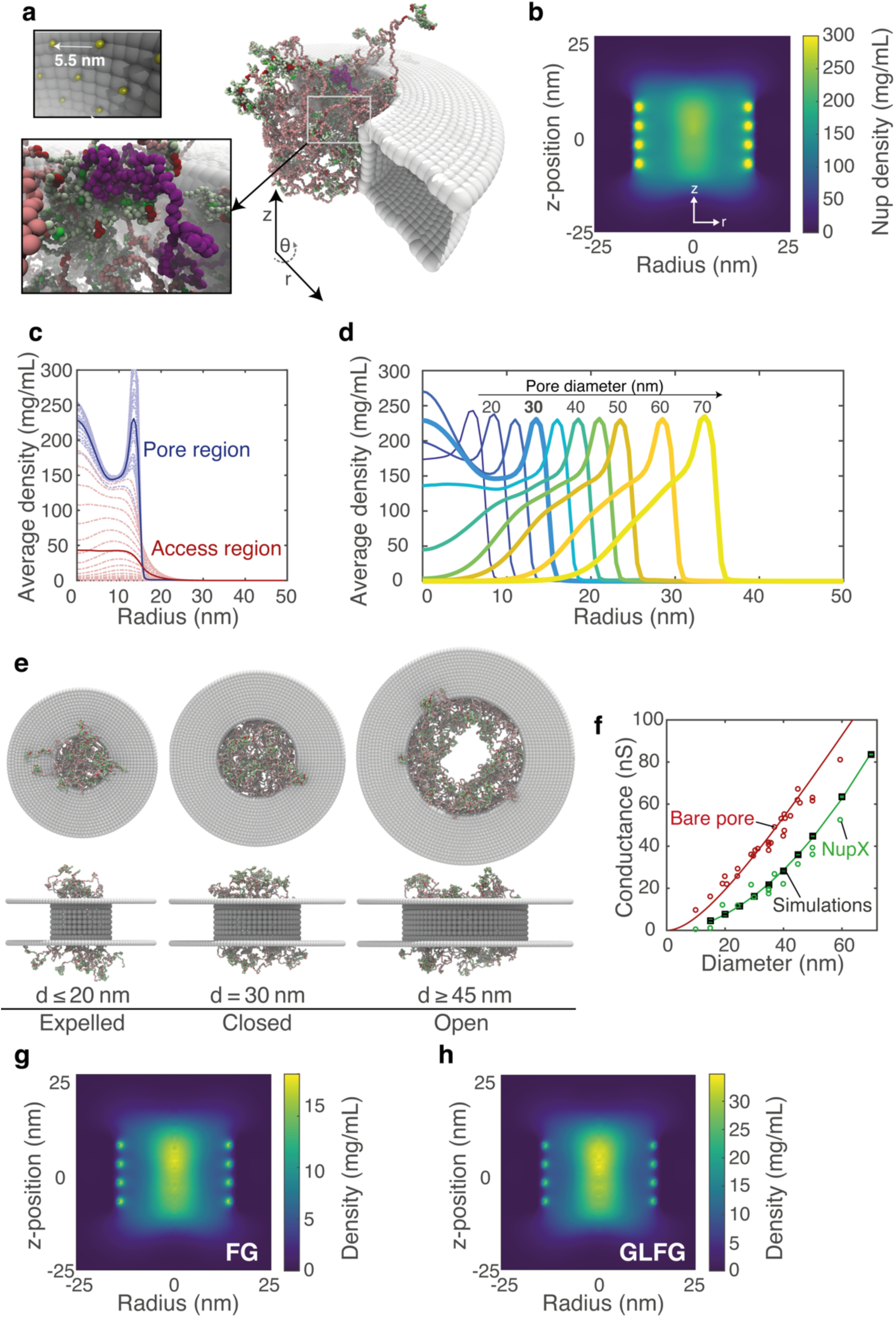
Protein distribution and conductance of NupX-coated pores. **a**, Snapshot of a biomimetic nanopore simulation. NupX proteins were tethered with a grafting distance of 5.5 nm (top inset) to a nanopore-shaped occlusion made of inert beads. Pore diameters ranged from 15-70 nm, where the pore thickness was 20 nm throughout. Bottom inset: highlight of a single NupX-protein (purple) within the NupX-meshwork. **b**, Axi-radial map (averaged over time and in the azimuthal direction) of the protein density within a 30 nm NupX-lined nanopore, from 6 μs simulations. The low C/H ratio domains of the NupX proteins form a high-density central ‘plug’. The high-density regions near the pore radius (15 nm) coincide with the anchoring sites of the NupX proteins. **c**, Density distributions for the pore (blue, |z| < 10 nm) and access (red, 10 nm < |z| < 40 nm) regions. Dashed curves indicate the average density within 1 nm thick slices in the z-direction. Thick curves indicate the z-averaged density profile for the pore and access regions, respectively. **d**, Radial density distributions (z-averaged) for NupX-lined nanopores with diameters ranging from 15 to 70 nm. The curve for 30 nm is emphasized. An increase in pore size beyond 30 nm leads to a decrease in the pore density along the pore’s central channel. **e**, Side-view and top-view visualizations of 20 nm, 30 nm, and 45 nm diameter NupX-lined nanopores. For nanopores with diameters smaller than 25 nm, the pore density decreases due to an expulsion of the collapsed NupX domains from the pore region towards the access region. For nanopores with diameters larger than 40 nm, the pore density decreases and a hole forms. For nanopores with a diameter of 25-30 nm, the pore region is sealed by the NupX cohesive domains. **f**, Conductance scaling for bare and NupX-coated nanopores. Open circles indicate conductance measurements for bare (red) and NupX-coated (green) pores. Squares indicate time-averaged conductance values obtained from MD simulations via a density-conductance relation (Materials and Methods). Error bars indicate the standard deviation in the time-averaged conductance and are typically smaller than the size of the marker. Second-order polynomial fits to the bare pore (experimental) and the simulated conductance values are included as a guide to the eye. **g,h**, Axi-radial density maps (averaged over time and in the azimuthal direction) for FG and GLFG motifs, respectively. Both types of motif localize in the dense central region, indicating that there is no spatial segregation of different types of FG-motifs in NupX-coated nanopores.

High-density regions form close to the attachment sites (i.e. the four dots at each wall in Fig. 4b) and along the central axis of the nanopore. From these data, we obtained a radial protein density profile, averaged over the pore height for the pore region (|*z*| < 10 nm, Figure 4c), which exhibits a maximum of 230 mg/mL at the pore center for the 30 nm NupX nanopore system. This density agrees well with values in the range of 200-300 mg/mL observed in earlier computational studies of the yeast NPC central channel^47,56^. We attribute the central localization of the NupX proteins to the combination of repulsion between the high C/H ratio extended domains near the pore wall and attraction between the cohesive, low C/H ratio collapsed domains of opposing NupX proteins. Since the average density in the ‘access region’ (10 nm < |*z*| < 40 nm, Figure 4c) is found to be low in comparison to the average density within the pore region, we conclude that the NupX proteins predominantly localize within the nanopore.

Upon decreasing the pore diameter to values below 25 nm, we find that the NupX collapsed domains are expelled from the pore region towards the access region, resulting in lower densities in the central pore region (Figures 4d,e). A dense central plug forms, however, for pore diameters between 25 and 35 nm. When the pore diameter is increased beyond 35 nm, an opening forms in the NupX meshwork near the pore centre (Figures 4d,e). This finding is consistent with the increased event frequency and translocation speed observed in 60 nm NupX-coated pores (Fig. S8). Using a relation between the local protein density and the local conductivity for the pore and access regions^31^, we calculated the conductance of the NupX nanopores for varying diameters (Figures 4d, S13, Materials and Methods). The calculated conductance from the simulated NupX-lined pores is shown in Figure 4f (black squares) together with the experimental conductances for bare and NupX-coated pores (open circles). An excellent correspondence is observed between the simulation and experimental results. Note that we adopted a critical protein density of 85 mg/ml from the earlier work on Nsp1^31^ while employing a different dependency of the conductivity on the local protein density (Materials and Methods). Interestingly, the slope of the conductance-diameter curve for NupX-lined pores converges to that of bare pores already at relatively small pore sizes. This is due to the formation of a hole within the NupX meshwork (Figure 4d) already in pores with diameters over 40 nm, rendering these effectively similar to bare nanopores of smaller diameter.

A spatial segregation of different types of FG-motifs, as was observed in recent computational studies^3,34^, is not studied here (Figures 4g,h). Instead, both types of FG-motifs localize similarly in the high-density central region within the NupX nanopore channel. From these distributions and the observed size-selective transport of these pores (Figures 3e-g), we can infer that a spatial segregation of different FG-motifs is not required for size-selective transport.

Finally, in order to assess the size-selective properties of NupX-lined nanopores, we simulated a 30 nm diameter NupX-lined nanopore in the presence of 10 Kap95 or 10 tCherry particles. We released Kap95/tCherry in the access region at the top and recorded their location in over 5 μs of simulation time (see Materials and Methods, Figs. 5c,d). The Kap95 particles entered and left the NupX meshwork and sampled the pore lumen by traversing in the z-direction (Fig. 5c). They localized preferentially at positions radially halfway between the central pore axis and the edge of the nanopore, where their time-averaged density distribution takes the shape of a concave cylindrical region, as is shown in Figure 5a. Kap95 was found to be capable of (re-)entering and leaving the meshwork on either side (Fig. 5c). Since no external electric field was applied, exiting and subsequent re-absorption of Kap95 into the NupX meshwork occurred and there was no directional preference for the motion of the Kap95 molecules, in contrast to the experiments. Interestingly, the NupX-meshwork adapted itself to the presence of the Kap95 particles by expanding towards the access region (compare Figs. 4b and S15): the protein density in the pore region decreased due to the presence of the Kap95, whereas the protein density increased in the access region. In contrast to the findings for Kap95, we observed that the sterically inert tCherry, simulated under the same conditions, remained in the top compartment (Fig. 5b,d) and did not permeate into the NupX-meshwork over the 5 μs time span of the simulation.

**Figure 5:**
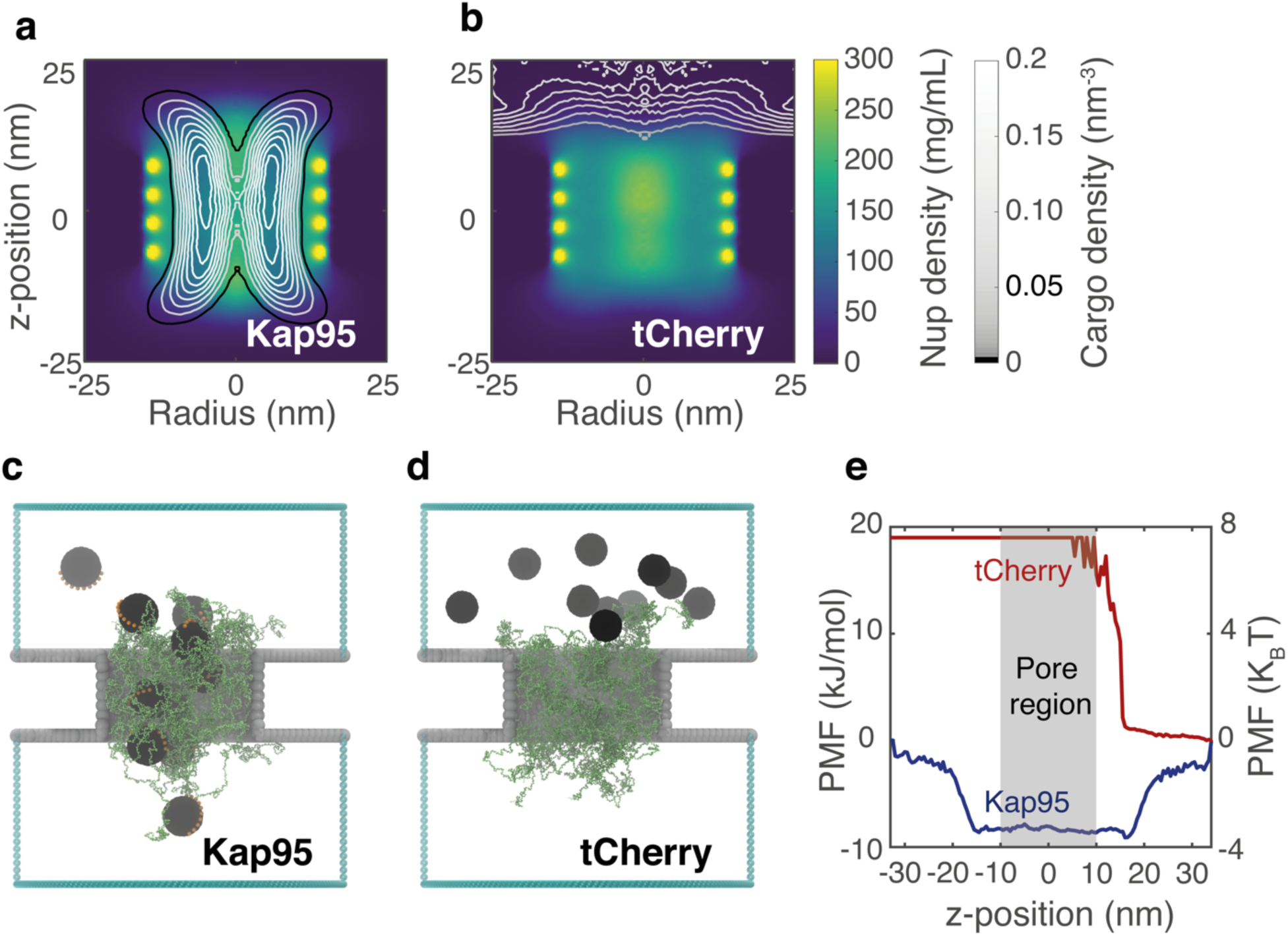
Effect of transporters on NupX-lined biomimetic pores. **a**, Contour graphs of the Kap95 number density (grey contours) superimposed on the NupX protein density distributions (in presence of Kap95) within a 30 nm NupX-lined nanopore (NupX-density is shown separately in Figure S15). The protein meshwork adapts (as compared to the distribution in Figure 4b) to accommodate the permeating Kap95 particles. **b**, Density distribution of tCherry superimposed on the NupX protein density distribution in a 30 nm diameter NupX-lined nanopore. tCherry remains in the top compartment and does not permeate the NupX-protein meshwork. **c, d**, Simulation snapshots of 30 nm NupX-lined nanopores in the presence of Kap95 particles (c, black spheres with orange binding spots) and tCherry particles (d, black spheres), which were released in the top compartment. Kap95 particles enter and exit the NupX meshworks at either side of the nanopore, whereas tCherry particles remain in the top compartment. **e**, PMF curves of Kap95 and tCherry along the z-coordinate, obtained via Boltzmann inversion of the normalized density profile along the z-axis. The pore region coincides with an energy well of over 3 *K*_*B*_*T* for Kap95, whereas tCherry experiences a steep energy barrier of ∼7 *K*_*B*_*T.*

To quantify the selectivity of the 30 nm NupX-lined nanopores, we calculated PMF-curves along the *z*-axis for both cargo types (Fig. 5e, see Materials and Methods). Kap95 experienced a negative free energy difference of approximately 8 kJ/mol, which amounts to a binding energy of just over 3*k*_*B*_*T* per Kap95. On the other hand, tCherry experiences a steep energy barrier of approximately 18 kJ/mol, which corresponds to over 7*k*_*B*_*T* per protein. From these results, we conclude that NupX-lined nanopores indeed reproduce the NPC’s remarkable selectivity towards Kap95.

## Discussion

In this work, we introduced a 311-residue long artificial FG-Nup, termed ‘NupX’, that we designed *de novo* based on the average properties of GLFG-type Nups (Nup49, Nup57, Nup100, Nup116, Nup145N). To our knowledge, the work presented here is the first demonstration of a fully artificial FG-Nup that, gratifyingly, was found to faithfully mimic the size-selective behaviour of the nuclear pore complex. We experimentally found that NupX-coated substrates bind selectively to Kap95 with a binding affinity of 191 ± 20 nM, while it did not interact with the control proteins BSA or tCherry – a finding confirmed through coarse-grained MD simulations of the absorption of Kap95 and tCherry particles. The binding affinity of Kap95 towards the NupX brush is very similar to that found in other *in vitro* studies on the interaction between human homologs of Kap95 and native FG-Nups^18,57^. Consistent with these results, we found that Kap95 translocates through both uncoated and NupX-lined nanopores on a physiological (∼ms) timescale^58^, whereas BSA passage through the NupX-coated pores was effectively excluded. Coarse-grained MD simulations revealed how the NupX proteins form a dense (>150 mg/mL) phase that allows passage of Kap95 particles while excluding inert particles. Interestingly, we find that the high densities of the FG-rich NupX meshworks are comparable to those obtained in earlier simulation studies of yeast NPCs^47^. A comparison of the intrinsic protein density (mass per unit Stokes volume) of NupX (219 mg/mL) with that of Nsp1 (74 mg/mL) explains why our NupX-meshworks localize more compactly inside nanopore channels than Nsp1 in earlier work^31^. The increased conductance of the denser NupX-lined nanopores (as compared to Nsp1) indicates that the average protein density is not the only factor that describes conductivity; the dynamics of the unfolded proteins and the local charge distribution might be important as well.

The design strategy presented in this work allows us to assess the role of the amino acid sequence of the spacer regions in GLFG-Nups. Spacer residues were reported to be involved in the interaction interface of Nup-NTR complexes^59,60,61^, highlighting a possible specific role of these domains in the binding of NTRs. In the current work, we assigned the positions of spacer residues along the NupX amino acid sequence entirely randomly, in both the collapsed, FG-rich low C/H ratio domain, and the extended high C/H ratio domain. This indicates that no specific spacer sequence motifs are required to facilitate the fast and selective transport of NTRs like Kap95. The consistency of the Stokes’ radii of different NupX designs within our simulations (Figure 1h) supports this finding.

Furthermore, our results shed light on the functional role of the spatial segregation of FG and GLFG motifs that was observed in earlier work^3,34^. Although these recent computational studies observed such a feature and suggested that it plays a role in size-selective transport, the coinciding distributions of FG and GLFG motifs (Figures 4g,h) show that no spatial segregation of different types of FG motifs exists within our size-selective nanopores. Notably, this does not rule out a different functional role for the spatial segregation of different types of FG-motifs, which can be explored in future work.

The combined design and characterization approach presented here, with brush-absorption and nanopore-transport measurements on the one hand and coarse-grained MD simulations on the other, provides a powerful and exciting platform for future studies of artificial FG-Nups: one can now start to systematically examine the relation between FG-Nup amino acid sequence and size selectivity of the NPC. Such studies could, for example, entail the design of FG-Nups with radically different physiochemical properties (i.e. FG-spacing, FG motif type, spacer domain C/H ratios, sequence complexity) to assess the size-selective properties of nanopore systems functionalized with these designer FG-Nups. Indeed, solid-state nanopores modified with a single type of FG-Nup were shown in this and other works^26,28,31^ to reproduce NPC size-selectivity, justifying the use of a single type of artificial Nups within an environment structurally similar to the NPC. Moreover, in view of the similarity^5^ and redundancy^21,35^ of different FG-Nups within the NPC and the ability of our method to robustly reproduce FG-Nup properties (Figures 1h and S11), we are confident that a single artificial FG-Nup can capture the complex transport behaviour of the NPC. However, given that even minimally viable native NPCs^21,35^ contain several different FG-Nups, it is worth mentioning that NPC mimics with a heterogeneous set of (artificial) FG-Nups can be created as well: DNA origami scaffolds^33^ potentially allow to position different artificial FG-Nups with great control, thus enabling systematic studies of how the interplay of different (artificial) FG-Nups gives rise to various transport properties of the NPC.

Finally, the design procedure that we introduced here is not limited to applications in nucleocytoplasmic transport. It may, for example, be possible to use a comparable approach to create *de novo* selective molecular filters (e.g. for use in artificial cells^62,63^), systems that would rely on selective partitioning of molecules in meshworks of unfolded proteins with assigned properties. Control can be asserted over the composition and geometry of the meshwork e.g. by means of recently developed DNA-origami scaffolds^32,33^. More generally, the approach illustrated here may enable future studies of the physical properties underlying phase separation of intrinsically disordered proteins^30^. One could, for example include degrees of freedom such as the proteins’ second virial coefficient (*B*_22_), or the charge patterning (*κ*), which have been linked to the phase behavior of intrinsically disordered proteins^64,65^. We envision that just like the field of *de novo* protein design has come to fruition with improved understanding of protein folding^66^, the design of unstructured proteins like NupX will enable a versatile platform to study the intriguing functionality of intrinsically disordered proteins.

## Acknowledgements

We would like to thank the Görlich lab for sharing purified Nsp1 and Nsp1-S proteins, and Jacklyn Novatt for the protocols on FG-Nup purification that we used for our artificial FG-Nup. We thank Meng-yue Wu for technical assistance on the TEM. This research was funded by NWO-I programme ‘Projectruimte’, grant no. 16PR3242-1. We acknowledge useful discussions with Adithya Ananth, Wayne Yang, Sergii Pud, Daniel Verschueren, and Sonja Schmid. H.W.d.V acknowledges support from the CIT of the University of Groningen and the Berendsen Centre for Multiscale Modeling for providing access to the Peregrine and Nieuwpoort high performance computing clusters. C.D. acknowledges support from the ERC Advanced Grant SynDiv (no. 669598) and the NanoFront and BaSyC programs.

## Data availability

Source data for Figs. 1d-h, 2a-i,l, 3c-g, 4b-d,f-h and 5a,b,e are provided with the paper in Supplementary Table S5. Other data that support the findings of this study are available from the corresponding authors upon reasonable request.

## Materials and Methods

### Analysis of GLFG-Nups and design of synthetic Nups

Protein sequences of *Saccharomyces cerevisiae* GLFG-type Nups (i.e. Nup100, Nup116, Nup49, Nup57 and Nup145N) were analyzed using a script custom-written with R programming package (version 3.3.1). Following the definitions of high C/H-ratio and low C/H-ratio unfolded FG-Nup domains as given in Ref. 5, we obtained histograms of the amino acid frequencies in both the collapsed (low C/H-ratio) and extended (high C/H-ratio) domains. The collapsed / extended domain sequences of NupX were then assigned in three steps: First, the collapsed and extended domains of NupX were assigned equal lengths of 150 residues each. Then, by normalizing the distributions in Figure 1d to the number of available residues within each domain, the total pool of amino acids within each domain was obtained. Lastly, these amino acids were randomly assigned a sequence index within each domain, with a boundary condition of the presence of FG and GLFG repeats spaced by 10 residues within the low C/H-ratio domain. This approach was repeated iteratively in combination with disorder predictions using the on-line PONDR disorder prediction utility^36^ until a sufficiently disordered design was obtained. The final version of the NupX amino acid sequence was also analyzed for secondary structure using DISOPRED^37,38^ and Phyre2^39^. A 6-histidine tag was added to the N-terminus of the NupX sequence in order to facilitate protein purification (see ‘Protein purification’ section). Finally, on the C-terminus a cysteine was included to allow the covalent coupling of the NupX protein to the surface.

### Expression and purification of NupX and Kap95

The synthetic NupX gene (Genscript), appended with codons for an N-terminal His6-tag and a C-terminal cysteine residue, was cloned into pET28a and expressed in Escherichia coli ER2566 cells (New England Biolabs, fhuA2 lacZ::T7 gene1 [lon] ompT gal sulA11 R(mcr73::miniTn10--TetS)2 [dcm] R(zgb-210::Tn10--TetS) endA1 δ(mcrCmrr)114::IS10). To minimize proteolysis of NupX, the cells were co-transformed with plasmid pED4, a pGEX-derivative encoding GST-3C-Kap95 under control of the tac promoter. Cells were cultured in shake flasks at 37 °C in Terrific Broth supplemented with 100 µg/mL ampicillin and 50 µg/mL kanamycin, and expression was induced at OD600∼0.6 with 1 mM IPTG. After 3 hours of expression the cells were harvested by centrifugation, washed with PBS, resuspended in buffer A1 (50 mM Tris/HCl pH 7.5, 300 mM NaCl, 8 M urea, 5 mg/mL 6-aminohexanoic acid supplemented with one tablet per 50 mL of EDTA-free cOmplete ULTRA protease inhibitor cocktail) and frozen as “nuggets” in liquid nitrogen. Cells were lysed with a SPEX cryogenic grinder, after thawing 1,6-hexanediol was added to a final percentage of 5%, and the lysate was centrifuged for 30 minutes at 40.000 rpm in a Ti45 rotor (Beckman Coulter). The supernatant was loaded onto a 5 mL Talon column mounted in an Akta Pure system, the column was washed with buffer A2 (50 mM Tris/HCl pH 7.5, 300 mM NaCl, 800 mM urea, 5 mg/mL 6-aminohexanoic acid, 2.5% 1,6-hexanediol) and NupX was eluted with a linear gradient of 0-200 mM imidazole. Peak fractions were pooled, diluted tenfold with buffer A2 lacking sodium chloride, loaded onto a 1 mL HiTrap SP sepharose HP column and NupX was eluted with a linear gradient of 0-1 M NaCl.

Kap95 was expressed as a C-terminal GST fusion protein in Escherichia coli ER2566 cells from plasmid pED4, a pGEX derived construct (kindly provided by Jaclyn Novatt) in which the thrombin cleavage site was replaced by a 3C protease cleavage site. Cells were grown in shake flasks at 30°C on LB medium supplemented with 100 µg/mL ampicillin, induction was induced at OD600∼0.6 with 1 mM IPTG, and growth was continued overnight. Cells were harvested by centrifugation, washed with PBS, resuspended in TBT buffer (20 mM HEPES/NaOH pH7.5, 110 mM KOAc, 2 mM MgCl2, 0.1% (w/v) Tween20, 10 µM CaCl2 and 1 mM β-mercaptoethanol), and lysed by a cell disruptor (Constant Systems) at 20 kpsi. Following centrifugation for 30 minutes at 40.000 rpm in a Ti45 rotor (Beckman Coulter), the supernatant was loaded onto a 2 mL GSTrap 4B column mounted in an Akta Pure system. The column was washed with TBT buffer, TBT + 1 M NaCl and TBT + 0.1 mM ATP, and the fusion protein was eluted with TBT + 10 mM reduced glutathione. The GST moiety was cleaved off by overnight digestion with home-made 3C protease, and Kap95 was separated from GST and the protease by size exclusion chromatography on a Superdex S200 column pre-equilibrated with TBT buffer.

### QCM-D sample preparation and data acquisition

QSense Analyzer gold- and SiN-coated quartz QCM-D chips were purchased from Biolin Scientific, Västra Frölunda, Sweden. Prior to the experiment, chips were immersed in RCA-1 solution, which consisted of 30% Ammonium Hydroxide, 30% Hydrogen Peroxide, and deionized (DI) water in 1:1:5 ratio, for ∼30 minutes at 75°C. This step was used to clean the surface from carbon species, as well as to enrich the surface with hydroxyl groups in case of the SiN-coated chips. Chips were further rinsed with DI water, sonicated for ∼10 minutes in pure ethanol, and blow-dried with a nitrogen stream. Before each experiment, flow-cells were disassembled, cleaned by sonication for 20-30 minutes in freshly prepared 2% SDS, rinsed with DI water, and blow-dried with a nitrogen stream. For SiN-coated quartz sensors, the SiN surface was chemically engineered in order to add free maleimide groups (see ‘Preparation of NupX-coated nanopores’ for details).

QMC-D data were monitored and recorded with sub-second resolution using Qsoft, which was provided by the company together with Qsense Analyzer. Buffer was injected into the flow-cell chamber at constant flow-rate of 20 μL/min using a syringe pump. Experiments were all performed at room temperature. Shift in the resonance frequency (δf) and dissipation (δD) can be, in first approximation, attributed to mass deposition and increase in viscoelasticity of the film, respectively. δf and δD were acquired at the fundamental tone (n = 1) and the 5 overtones (n = 3, 5, 7, 9, 11). The normalized second overtone 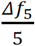 was used for display and analysis. Data processing, filtering, and plotting were performed using a custom-written Matlab script.

### Fitting of the QCM-D data with a viscoelastic model and estimation of the grafting density

QCM-D were fitted using the Voigt-Voinova viscoelastic model^40,42^ to extract the adsorbed areal mass density m_Voigt_, using the Qtools software (provided by Biolin Scientific), which represents the ‘wet mass’ as it includes both the ‘dry mass’ of the protein m_protein_ as well as the hydrodynamically coupled water m_water_. For the modelling, we assumed as input parameters a fluid density of 1000 kg/m^3^, a fluid viscosity of 0.001 kg/ms, and a film density of 1 g/cm^3^, as typically used in soft matter^41^. The harmonics employed for the fitting were the 3^rd^, 5^th^, and 7^th^ for both frequency and dissipation. The protein mass was estimated by assuming that 35% of m_Voigt_ is constituted by m_protein_, while the remaining 65% is coupled water m_water_, consistent with previous work on combining QCM-D with optical techniques^43^.

### Preparation of NupX-coated nanopores and current data acquisition

Solid-state nanopores with diameters from 10-60 nm were drilled using TEM in glass-supported SiN_x_ free-standing membranes. Glass chips were purchased from Goeppert. We refer to Ref. 46 and 58 for details on the fabrication of the chip substrate and free-standing membrane. Freshly drilled solid-state nanopores were rinsed with ultrapure water, ethanol, acetone, and isopropanol, followed by 2-5 minutes of oxygen plasma treatment, which was performed in order to further clean and activate the nanopore surface with hydroxyl groups. Next, chips were incubated in 2% APTES (3-aminopropyl-triethoxysilane) (Sigma Aldrich) in anhydrous toluene (Alfa Aesar) for 45-60 min at room temperature, shaking at 400 rpm, followed by 15 minutes in anhydrous toluene for washing. These two steps were performed in a glove-box under constant nitrogen stream in order to prevent the APTES from polymerizing. Then, chips were further rinsed with ultrapure water, ethanol, and heated at 110°C for at least 30 minutes. This step was used to fixate the APTES layer by favouring further binding between the unreacted ethoxy groups.

The nanopore surface was thus covered with primary amines, which were subsequently reacted to Sulfo-SMCC (sulphosuccinimidyl-4-(N-maleimidomethyl)-cyclohexane-1-carboxylate) (2 mg no-weight capsules (Pierce)), a crosslinker that contains NHS-ester (reacts to amines) and maleimide (reacts to thiols) groups at opposite ends, for > 3hrs at room temperature, shaking at 400 rpm. Chips were subsequently washed in PBS for 15 minutes and incubated with thiolated proteins for 2-3 hours, which were pretreated with 5 mM TCEP for ∼30 minutes in order to reduce the thiol groups. Chips were further washed in PBS before the electrical measurement. Raw ionic current traces were recorded at 100 kHz bandwidth with an Axopatch 200B (Molecular devices) amplifier, and digitized (Digidata 1322A DAQ) at 250 kHz. Traces were monitored in real-time using Clampex software (Molecular devices). Data were digitally filtered at 5 kHz using a Gaussian low-pass filter and analysed using a custom-written Matlab script^68^.

### Dynamic light scattering (DLS) measurement of the hydrodynamic diameter

DLS experiments were performed using Zetasizer Nano ZS (Malvern). Cuvette of 100 μL (Brand GMBH) were used for the measurement. All protein hydrodynamic diameters were measured in 150 mM KCl, 10 mM Tris, 1mM EDTA, at pH 7.5, and averaged over three experiments. Mean value and standard deviation for each of the proteins used are reported in Table S1. Proteins which contained exposed cysteines (NupX, Nsp1, and Nsp1-S) were pre-treated with TCEP (present in at least 100x excess) in order to break disulfide bonds.

### Coarse-grained model for unfolded proteins

All coarse-grained MD-simulations were performed using our earlier developed one-bead-per-amino acid (1BPA) model for unfolded proteins^47,69^. This model maps complete amino acids to single beads with a mass of 124 amu placed on the Cα position, separated by an average bond length (modelled as a stiff harmonic potential) with an equilibrium distance of 0.38 nm. Backbone potentials were assigned via an explicit coarse-grained mapping of Ramachandran data of a library of the coil regions of proteins that distinguishes flexible (i.e. Glycine), stiff (i.e. Proline) and regular amino acids^69^. Non-bonded interactions between different amino acid residues are based on their respective hydrophobicity (normalized between 0 and 1 and based on the free energy of transfer between polar and apolar solvents) and obey the following interaction potential^47^:

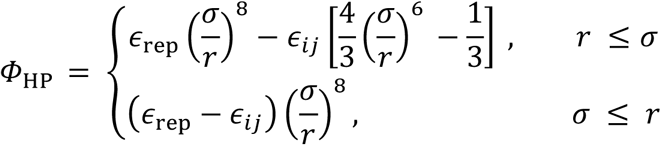

where *ϵ*_rep_=10 kJ/mol and 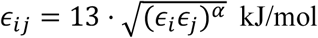, with *ϵ*_*i*_ the normalized (between 0 and 1) hydrophobicity of a residue *i* and *α* = 0.27 a scaling exponent. The electrostatic interactions within the 1BPA model are described by a modified Coulomb law:

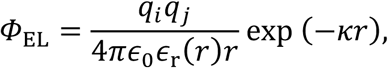

where the electrostatic interactions are modulated via a Debye screening component. This form of electrostatics takes into account the salt concentration (set at 150 mM here, via a screening length *κ* =1.27 nm^-1^) together with a solvent polarity at short distances via a distance-dependent dielectric constant:

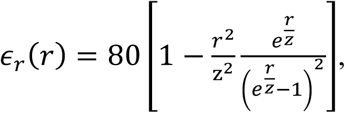

where *z* = 0.25. Non-bonded interactions are cut-off at 2.5 nm (hydrophobic interactions) or 5.0 nm (electrostatic interactions). Since the 1BPA model operates without explicit solvent, we apply stochastic dynamics with a coupling frequency *τ*_*T*_ of 50 ps^-1^. Stochastic dynamics handles temperature coupling implicitly, ensuring that the system operates within a canonical ensemble at a reference temperature of 300 K. We refer the reader to the original work for further details on the used 1BPA model^47,69^. Unless otherwise mentioned, all simulations were performed using the above forcefield and corresponding settings, employing the GROMACS^70^ molecular dynamics software (version 2016.1/2016.3) on a parallelized computer cluster. A complete overview of all simulations in this work is provided in Table S3.

### Calculating Stokes radius of NupX and variations

Intrinsically disordered proteins were modelled using the 1BPA model^47,69^, starting from an extended configuration. After energy minimization (steepest descent) and a brief (5 ns) equilibration step, we simulated the individual proteins for 5 · 10^8^ steps using a timestep of 20 fs (total simulation time: 10 μs). Conformations were extracted every 10000 frames (i.e., every 200 ps). In order to calculate the Stokes radii (*R*_*S*_) from the MD trajectories, we extracted protein conformations every 2 ns and applied the HYDRO++ software^71^ in order to calculate the *R*_*S*_-values. This procedure yields a total of 5000 Stokes radii per protein.

### Calculating NupX brush density profiles and PMF curves of cargo absorption

We modelled the brush substrate as a fully triangulated (sometimes denoted as ‘hexagonally’) closed-packed array of sterically inert beads with a diameter of 3 nm. NupX proteins were tethered on top of the scaffold by their C-terminus in the densest way possible, which is on an equilateral triangular lattice with an average grafting distance of 5.7 nm. A fully triangulated lattice is close-packed in two dimensions, meaning that a unique length scale sets the grafting density. The simulation box consisted of a 34.2×34.2×81.5 nm^3^ triclinic and fully periodic unit cell (Figs. 2j and k). The grafting pattern of the NupX proteins was placed such as to ensure homogeneity of the NupX-brush in the lateral plane throughout the periodic boundaries. Density profiles for the NupX brushes and FG/GLFG repeats were obtained by simulating the NupX brush systems for 1.75 × 10^8^ steps (3.5 μs) using a timestep of 20 fs. The first 500 ns of the simulation trajectory was discarded as equilibration. We modelled Kap95 and tCherry in the following way: The Kap95 particle consists of sterically repulsive beads, arranged in a geodesic shell such that the particle has a diameter of 8.5 nm, consistent with the hydrodynamic dimensions of the Kap95 protein (Table S1). The Kap95 surface beads interact with the NupX amino acid beads through the repulsive term of *ϕ*_*EL*_ (i.e., volume exclusion), and the modified Coulomb potential *ϕ*_HP_, where we distributed the charge (total net charge of −43e) of Kap95 over the Kap95 surface beads. We preserved the structure of the particle by applying a harmonic restraint of 40000 kJ/mol on bead pairs whenever the distance between beads within the reference structure was below a cut-off of 1 nm. A total of ten hydrophobic binding pockets were placed at a mutual distance of 1.3 nm along an arc (Figure S14) on the surface of the Kap95-particle^8,31,48,49^. The binding sites interact with NupX amino acid beads via the hydrophobic potential *ϕ*_HP_, where the hydrophobicity of these binding sites was set equal to that of Phenylalanine. In the same way as for the Kap95, we assembled a tCherry-sized particle^10,31,49^ with a diameter of 7.5 nm (Table S1). Other than steric repulsion, no specific interactions between tCherry on one hand, and the amino acid or substrate beads on the other were assigned. Using a harmonic restraint of 100 kJ/mol, we generated umbrella sampling windows by pulling the cargo in the negative z-direction (while freezing the particle’s movement in the *xy*-plane) with a pulling velocity of -0.001 nm/ps and a time step of 20 fs along the centre of the triclinic box. Starting configurations were extracted every 0.5 nm, yielding 51 umbrella windows per cargo. After energy minimization (removal of overlap between beads) via the steepest descent algorithm, we performed 100 ns (5×10^6^ steps) of equilibration, and 1 μs (5×10^7^ steps) of production MD per umbrella window, where the Kap particles were restrained using a harmonic umbrella potential of 100 kJ/mol in the z-direction, applied to the cargo’s center of mass. Aside from this restraint, the particles were free to rotate and move in the *xy*-plane. Potential-of-mean-force (PMF) curves were obtained using the weighted histogram analysis method (WHAM) via the g_wham^50^ utility of GROMACS.

### Coarse-grained MD simulations of NupX-lined nanopores

We modelled the SiN nanopores as cylindrically shaped occlusions in a membrane constituted entirely of sterically repulsive beads with a diameter of 3 nm. The height of the nanopore was 20 nm in all cases, with diameters ranging from 15 to 70 nm. NupX-proteins were modelled using the 1BPA model described earlier and tethered to the inner surface of the cylinder in an equilateral triangular lattice with a grafting distance of 5.5 nm. This value was estimated from NupX immobilization on SiN surfaces as measured through QCM-D and is in agreement with estimates used in earlier work^31^. Simulations were carried out for 2×10^8^ steps using a timestep of 15 fs (3 μs), or 4×10^8^ steps (6 μs) for the single case of a 30 nm diameter NupX-lined nanopore.

### Density distributions and nanopore conductance from nanopore simulations

Axi-radial density maps were obtained from NupX nanopore simulation trajectories using the ‘gmx_densmap’ utility of GROMACS, where a bin size of 0.5 nm was used to construct number densities within a cylinder centred on the nanopore. Average densities were extracted for the ‘pore’ and ‘access’ regions by averaging the axi-radial density distributions over the coordinate ranges |*z*| ≤ 10 nm and 10 nm < |*z*| < 40 nm, respectively^31^.

The conductance of NupX-lined nanopores was obtained by assuming that the conductance *G(d)* is governed by a modified Hall-formula^31,53^:

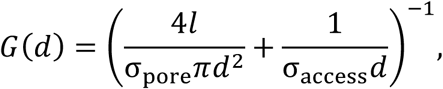

where *l* = 20 nm is the height of the nanopore, *d* denotes the diameter (15-70 nm), and *σ*_pore_ and *σ*_access_ denote the conductivities in the pore and access regions, respectively. The conductivities in both regions can be extracted from the axi-radial density distributions by integrating and normalizing the local conductivity over the pore diameter and corresponding height ranges:

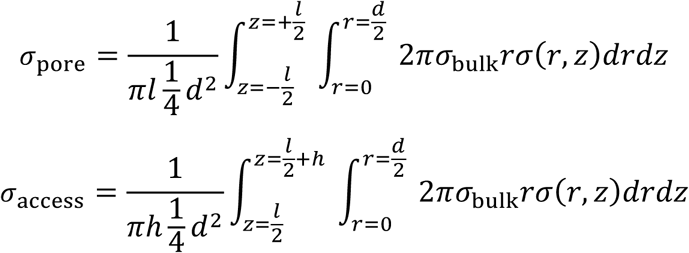

The local conductivity *σ*(*r, z*) follows from the local axi-radial density distribution *ρ*(*r, z*):

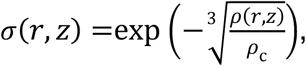

where *ρ*_*c*_ is set to 85 mg/mL^31^. Whereas this relation is a zero-parameter fit, the dependency of the conductivity on the local protein density is different from the linear model with a strict cut-off used in earlier work^31^. Here, the conductivity drops quickly with protein density while decreasing only slowly at high protein densities. This change in dependence was necessary since NupX-lined pores show a higher conductance at higher densities than the NPC mimics in earlier work. Axi-radial density distributions and the corresponding conductance were calculated for 100 ns windows to obtain an average conductance for each pore size. The sensitivity of this method was tested against the time averaging window of the density distributions and was found to only be marginally influenced by the window size, nor did the conductance change with time.

### Size selectivity of NupX-lined nanopores

We probed the size-selectivity of a NupX-lined nanopore with a diameter of 30 nm by inserting either 10 Kap95 molecules or 10 tCherry molecules to the *cis*-side of the nanopore. To speed up sampling, Kap95 or tCherry particles were confined to the periphery of the nanopore using a cylindrical constraint that prevents Kap95 or tCherry from entering regions with |*z*| > 40 nm or *r* > 35 nm^72^. This occlusion consisted entirely of sterically repulsive beads with a diameter of 3 nm, which only interact with Kap95 or tCherry. The size of the cylindrical constraint was set such that Kap95 or tCherry molecules can only leave the access region by entering the NupX-lined nanopore. The total simulation time spanned 3.33×10^8^ steps (5 μs), where we discarded the first 10% (500 ns) of the trajectory as equilibration. Axi-radial density maps for the protein density and contour graphs for the Kap95 and tCherry densities were obtained using the gmx_densmap utility using a bin size of 0.5 nm. We reported a cargo number density in lieu of a mass density, since the number of beads that constitute the Kap95 or tCherry-sized particle does not necessarily correspond to the amount of amino acids in either protein.

To calculate the PMF curve along the *z*-axis for Kap95 and tCherry, we integrated the axi-radial number density of the centre-of-mass of the particles in the radial direction, resulting in a one-dimensional axial number density *n*(*z*). Normalizing *n*(*z*) with the number of bins in the *z*-direction results in a probability distribution *p*(*z*), from which the PMF can be calculated by using the inverse Boltzmann relation:

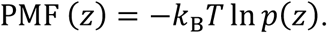

The PMF curves for both Kap95 and tCherry (Fig. 5e) were shifted such that the curves were zero at *z* = 35 nm. Regions with zero density (leading to divergence of the ln *p*(*z*)-term) were set equal to the maximum of the PMF curve.

